# Export‑biased, 3′UTR‑preserving TDP‑43 model links nuclear loss to cytoplasmic aggregation in ALS/FTLD

**DOI:** 10.1101/2025.11.12.687969

**Authors:** Genri Toyama, Shingo Koide, Takuma Yamagishi, Aya Washida, Ryutaro Hanyu, Osamu Onodera, Akihiro Sugai

**Author notes:** Corresponding author. Email: Akihiro Sugai.

## Abstract

Nuclear depletion and cytoplasmic mislocalization of TDP-43 are central pathological features of amyotrophic lateral sclerosis and frontotemporal lobar degeneration. TDP-43 protein levels are normally maintained by autoregulation through its native 3′ untranslated region (3′ UTR). It remains untested whether this feedback is still protective once the perturbation is chronic, and whether the resulting rise in *TARDBP* transcripts restores the functional nuclear pool or is instead diverted into non-functional species. To address this, we engineered full-length human TDP-43 carrying an N-terminal nuclear export signal (NES) while leaving the native 3′ UTR autoregulatory module intact. Expression was titrated so that whole-cell RIPA-soluble exogenous TDP-43 remained ≤30% of endogenous levels. In HEK293T cells, export bias drove TDP-43 into the cytoplasm and produced detergent-insoluble species. In differentiated SH-SY5Y cells, nuclear splicing defects and autoregulatory changes scaled with export-biased load, and detergent-insoluble accumulation was already present within the same low-load range. Human iPSC-derived neurons showed a comparable cytoplasmic shift, discrete TDP-43-positive foci, and TDP-43-dependent splicing defects. When endogenous TDP-43 was selectively depleted under transgene induction, 3′ UTR-coupled autorepression was weakened and transgene-derived *TARDBP* transcripts rose; however, the additional exogenous TDP-43 did not expand the soluble, splice-competent pool but partitioned into insoluble fractions. Splicing defects in differentiated SH-SY5Y cells persisted even when soluble NES–TDP-43 reached endogenous-equivalent levels. We therefore propose that under native 3′ UTR control, sustained cytoplasmic bias uncouples compensatory *TARDBP* upregulation from recovery of the functional nuclear pool; the extra output is diverted into insoluble, fragmented species rather than restoring nuclear function.

## 1. Introduction

TDP-43 is an essential RNA-binding protein whose abundance must be held within a narrow functional range (Lee et al., 2011; Ling et al., 2013; Sugai et al., 2018; Tziortzouda et al., 2021). This homeostasis depends on coordinated nucleocytoplasmic transport, maintenance of protein solubility, and autoregulation of *TARDBP* expression through the native terminal exon/3′ untranslated region (3′ UTR) (Avendaño-Vázquez et al., 2012; Johnson et al., 2009; Walker et al., 2015). In amyotrophic lateral sclerosis (ALS) and frontotemporal lobar degeneration (FTLD), the affected neurons typically show both nuclear depletion and cytoplasmic accumulation of TDP-43 (Arai et al., 2006; Neumann et al., 2006). How these two features become coupled, and how autoregulatory feedback shapes that coupling, has received relatively little direct attention.

The native 3′ UTR-coupled circuit adjusts *TARDBP* output in response to changes in functional nuclear TDP-43 (Ayala et al., 2011; Polymenidou et al., 2011; Tollervey et al., 2011), and a transient drop in nuclear function should therefore be buffered. Consistent with this, mislocalization or aggregation of TDP-43 has been shown to elicit compensatory *TARDBP* upregulation alongside nuclear loss-of-function phenotypes (Koyama et al., 2016; Mamede et al., 2026). What is not yet clear is whether the same buffering still works when the perturbation does not resolve. If the additional *TARDBP* output reaches the nucleus as functional protein, homeostasis is restored; if it is consumed instead by insoluble or fragmented cytoplasmic species, the compensatory response itself contributes to disease.

We addressed this by engineering an export-biased model of full-length human TDP-43 that retains the native terminal exon/3′ UTR autoregulatory module, so that the consequences of a chronic cytoplasmic shift could be examined while preserving RNA-level feedback. We then combined biochemical, localization, and splicing readouts across HEK293T cells, differentiated SH-SY5Y cells, and human iPSC-derived neurons to ask where the compensatory *TARDBP* output goes under persistent cytoplasmic bias—back into the functional nuclear pool, or into insoluble and fragmented cytoplasmic species.

## 2. Methods

### 2.1. Experimental design and rationale

To examine the consequences of persistent cytoplasmic bias under native autoregulatory control, we used a full-length NES–TDP-43 construct retaining the native terminal exon/3′ UTR (Fig. 1A). We defined low-load condition as whole-cell RIPA-soluble exogenous TDP-43 ≤30% of endogenous levels; this threshold was chosen to limit overexpression artifacts and reflects whole-cell soluble abundance, not physiological compartment-specific concentrations. Three readouts were predefined: (i) increased detergent-insoluble TDP-43 (RIPA-insoluble/urea-soluble) and C-terminal fragments; (ii) reduced nuclear splicing competence, measured by *STMN2* cryptic-exon inclusion and *POLDIP3* exon-3 skipping (Melamed et al., 2019; Shiga et al., 2012); and (iii) weakened 3′ UTR-mediated autorepression, quantified as a decrease in the exitron exclusion-to-inclusion ratio (Ex/In). The *TARDBP* exitron is an exon-embedded spliceable element (previously termed intron 6; Ayala et al., 2011; Koyama et al., 2016). A decrease in Ex/In indicates a shift toward the stable, exitron-included transcript consistent with reduced autorepression.

**Figure 1.**
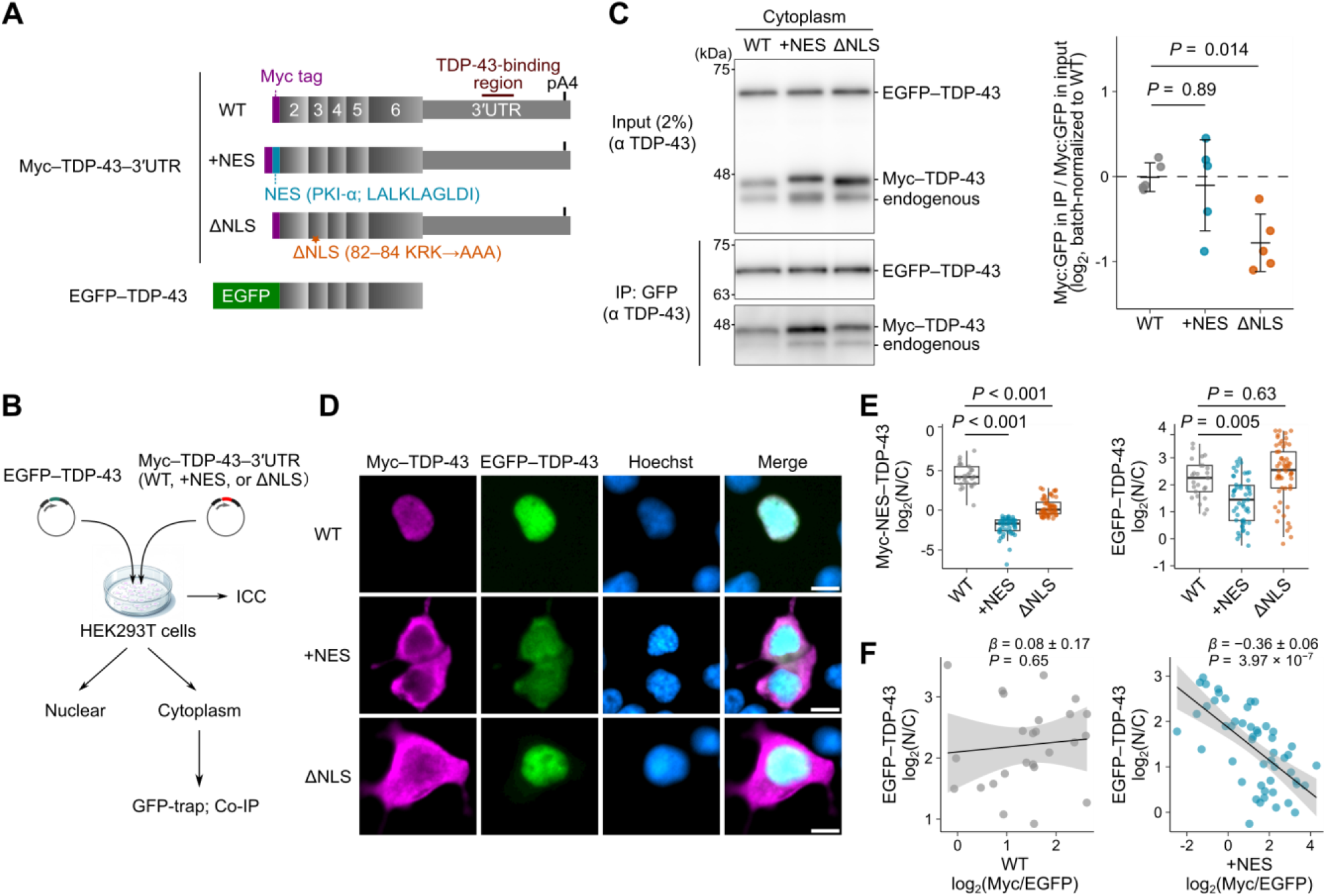
NES–TDP-43 preserves NTD-mediated dimerization and redistributes co-expressed WT TDP-43 toward the cytoplasm. (A) Construct schematics. Full-length human TDP-43 variants (WT, +NES, and ΔNLS) were N-terminally Myc-tagged and retained the full-length 3′ UTR. EGFP–TDP-43 lacking the native 3′ UTR served as the bait for interaction assays. (B) Experimental workflow. HEK293T cells co-expressing EGFP–TDP-43 and Myc–TDP-43 variants were processed for immunocytochemistry or cytoplasmic GFP-Trap co-immunoprecipitation (Co-IP). (C) Cytoplasmic Co-IP under RNase-free conditions. Immunoblots of cytoplasmic inputs and GFP pulldowns (IP). Association index = log_2_[(Myc/EGFP)_IP / (Myc/EGFP)_Input], normalized to WT. One-way ANOVA (*F*(2, 12) = 6.18, *P* = 0.014, *n* = 5) followed by Dunnett’s post hoc test versus WT. Data represent mean ± SD. (D) Representative images showing Myc–TDP-43 (magenta) and EGFP–TDP-43 (green). Scale bars, 10 µm. (E) Quantification of nucleocytoplasmic ratio (N/C) for each Myc-tagged variant and EGFP–TDP-43 (*n* = 132 cells from 28 image fields). Linear mixed-effects model (LMM) with image field as random intercept and Dunnett’s contrasts versus WT. (F) Single-cell co-recruitment analysis. EGFP–TDP-43 localization [log_2_(N/C)] plotted against +NES driver stoichiometry [log_2_(Myc/EGFP)]. Line indicates fitted regression; shaded band, 95% CI (LMM regression).

### 2.2. Plasmids and cloning

We used a tetracycline-inducible PiggyBac vector expressing N-terminally Myc-tagged human *TARDBP* as the backbone (VectorBuilder, VB241128-1134yxe). The *TARDBP* coding sequence (NM_007375.4) was fused to the native full-length 3′ UTR to generate a 3′ UTR-preserving construct (Fig. 1A; WT). From this backbone, we generated +NES by inserting the PKIα-derived nuclear export signal peptide (LALKLAGLDI) (Wen et al., 1995) immediately downstream of the Myc tag, and ΔNLS by mutating the nuclear localization signal (NLS) (KRK^82–84^) to AAA^82–84^ by site-directed mutagenesis (Barmada et al., 2010; Winton et al., 2008); ΔNLS served as a mislocalization control. For transient co-recruitment assays, EGFP–TDP-43 lacking the native 3′ UTR (VectorBuilder, VB240423-1225beg) was used as the bait construct. All constructs were verified by Sanger sequencing.

### 2.3. Cell models, expression paradigms, and induction

HEK293T cells (RRID: CVCL_0063; ECACC) were STR-authenticated, confirmed mycoplasma-negative (Biological Industries Ltd., 20-700-20), and maintained in DMEM supplemented with 10% FBS. For transient co-expression assays, cells were co-transfected with EGFP–TDP-43 and Myc-tagged variants using X-tremeGENE HP DNA transfection reagent (Roche, XTGHP-RO) and harvested 48 h later. The plasmid mass ratio was set to EGFP–TDP-43:Myc-variant = 1:20 to approximate comparable protein levels.

To establish an inducible export-biased system, a single-clone Tet-On HEK293T line expressing *Myc-NES– TDP-43–3′ UTR* was generated using a hyperactive PiggyBac transposase (mCherry-hypb; VectorBuilder, VB160216-10057) and maintained in DMEM supplemented with 10% tetracycline-free FBS (Tet System Approved FBS; Takara Bio, Z1106N). Doxycycline titration was used to define the low-load whole-cell soluble window described above; unless otherwise specified, cells were induced with 1.0 µg mL^−1^ doxycycline for 48 h.

To examine 3′ UTR-coupled feedback under endogenous depletion while holding imposed export bias constant, Tet-On cells were induced with doxycycline (1.0 µg mL^−1^) and concurrently transfected with 20 nM siRNA targeting the *TARDBP* 5′ UTR (5′-CGGCCTAGCGGGAAAAGTA-3′; Dharmacon) or a non-targeting control siRNA (IDT, 51-01-14-04) using Lipofectamine RNAiMAX (Thermo Fisher Scientific, 13778075). Cells were analyzed 72 h after siRNA transfection.

### 2.4. Neuronal differentiation and lentiviral transduction

SH-SY5Y cells (RRID: CVCL_0019) were maintained in DMEM/F12 GlutaMAX (Gibco, 10565018) supplemented with 10% FBS. For neuronal differentiation, cells were plated on Matrigel-coated plates (Corning, 356231) and cultured from day 0 in low-serum medium (1% FBS) supplemented with 10 µM all-trans retinoic acid (ATRA) (FUJIFILM Wako, 186-01114) and 50 ng mL^−1^ brain-derived neurotrophic factor (BDNF) (FUJIFILM Wako, 028-16451), with half-medium changes every 2–3 days. On day 5, cells were transduced with lentivirus encoding *Myc-NES–TDP-43–3′ UTR* (VectorBuilder, VB2503221105egs) or mCherry as a transduction control (VB2506191277hez) at varying multiplicities of infection (MOI; range 1.5– 4.5 for low-load conditions, MOI 10 for soluble-matched conditions). To increase effective viral concentration, half of the culture medium was removed immediately before viral addition and replaced with fresh medium 24 h later. Differentiation was verified by neuron-like morphology and βIII-tubulin (TUJ1) immunoreactivity.

Commercially available human iPSC-derived neurons (ReproNeuro; REPROCELL RCDN001N) were seeded onto ReproNeuro Coat (RCDN201)-coated plates and maintained in ReproNeuro Culture Medium (RCDN101) according to the manufacturer’s instructions. Half-medium changes were performed on days 3 and 7. Cells were transduced on day 14 *in vitro* (DIV14) as described for SH-SY5Y cells. Transduced neurons were analyzed at DIV21.

### 2.5. Subcellular fractionation and solubility assays

Nucleocytoplasmic fractionation was performed using NE-PER reagents (Thermo Fisher Scientific, 78833) supplemented with protease inhibitor cocktail (PIC), according to the manufacturer’s instructions. Fraction purity was assessed using Lamin B1 as a nuclear marker and GAPDH as a cytoplasmic marker. To quantify detergent-insoluble material from the same samples, the residual pellet was washed once in RIPA buffer (FUJIFILM, 188-02453; 50 mM Tris-HCl pH 8.0, 150 mM NaCl, 1% NP-40, 0.5% sodium deoxycholate, 0.1% SDS) containing PIC, sonicated on ice (20% amplitude, 10 s, two cycles), and ultracentrifuged (100,000 × *g*, 30 min, 4°C). The resulting pellet was solubilized in urea buffer (7 M urea, 2 M thiourea, 4% CHAPS, 30 mM Tris pH 8.5) containing PIC, sonicated, and centrifuged (15,000 × *g*, 20 min, 22°C). The supernatant was collected as the RIPA-insoluble (urea-soluble) fraction.

For whole-cell solubility profiling, cells were lysed directly in RIPA buffer containing PIC, sonicated, and ultracentrifuged (100,000 × *g*, 30 min, 4°C). The supernatant was collected as the RIPA-soluble fraction. The RIPA-insoluble pellet was re-extracted in urea buffer as above.

RIPA-soluble protein concentrations were measured by BCA assay (Thermo Fisher Scientific, 23225). Urea fractions were loaded at a fixed proportion relative to their paired RIPA-soluble lysates to preserve matched soluble/insoluble comparisons. Proteins were resolved by SDS–PAGE (10% SuperSep Ace), transferred to PVDF membranes, and probed with anti-TDP-43 (Proteintech, 12892-1-AP or Cosmo Bio, TIP-TD-P09), anti-Myc (9B11; Cell Signaling Technology, 2276S), anti-Lamin B1 (MBL, PM064), and anti-GAPDH (MBL, M171-3). Chemiluminescence (Millipore, WBKLS) was captured using an Amersham Imager 680. Densitometry of RIPA-soluble fractions was normalized lane-wise using total protein detected with No-Stain Protein Labeling Reagent (Invitrogen, A44717); urea-soluble fractions were normalized to their paired RIPA-soluble loading controls.

### 2.6. Protein interaction and co-recruitment assays

Co-immunoprecipitation was performed using the GFP-Trap Magnetic Agarose kit (Proteintech, GTMAK-20). HEK293T cells co-expressing EGFP–TDP-43 and Myc-tagged variants were lysed in non-denaturing buffer (14 mM Tris-HCl pH 7.5, 15 mM NaCl, 0.1% NP-40, 0.5 mM EDTA, protease inhibitors), clarified by centrifugation (17,000 × *g*, 10 min, 4°C), and adjusted to the manufacturer-recommended binding conditions. To distinguish predominantly protein-mediated association from RNA-bridged interactions, parallel lysates were treated with RNase A (40 µg mL^−1^; FUJIFILM Wako) and RNase T1 (100 U mL^−1^; Takara Bio) for 20 min before pulldown. Association efficiency was defined as (Myc/EGFP signal in the IP)/(Myc/EGFP signal in the input).

For image-based co-recruitment analysis, fluorescence images were segmented to define nuclear (Hoechst) and cytoplasmic compartments. For each cell, the nuclear-to-cytoplasmic ratio (N/C) of EGFP–TDP-43 was calculated. To separate redistribution effects from variation in driver expression, EGFP–TDP-43 localization [log_2_(N/C)] was regressed against driver stoichiometry [log_2_(Myc/EGFP)]. The slope of this relationship was used as a measure of dose-dependent redistribution of EGFP–TDP-43 as a function of export-biased driver abundance.

### 2.7. RNA isolation and splicing analysis

Total RNA was extracted using the NucleoSpin RNA kit (Takara Bio) with on-column DNase digestion and reverse-transcribed using PrimeScript RT Master Mix (Takara Bio). qPCR was performed with TB Green chemistry (Takara Bio) on a Thermal Cycler Dice System III. To assess *TARDBP* exitron processing as a readout of autoregulatory state, we quantified the two dominant exitron-excluded isoforms (*TARDBP* c.842– 1783 and c.833–1783, generated from alternative 5′ splice sites within the *TARDBP* exitron) together with the exitron-included isoform. Isoform abundance was normalized to *RPLP1* and reported as log_2_ fold-change. For summary analyses, Ex was defined as the sum of the two dominant exitron-excluded isoforms and In as the exitron-included isoform; autoregulatory state was summarized as log_2_(Ex/In).

Because isoform-specific RT-qPCR in +NES conditions measures pooled endogenous and transgene-derived *TARDBP* transcripts, pooled Ex/In was interpreted together with isoform-level deconvolution and source-specific assays. The splice-competent transgene contributes to the exitron-excluded pools; so a net decrease in exitron-excluded signal in pooled assays is compatible with reduced endogenous exitron exclusion. To distinguish transcript origin, endogenous transcripts were detected with primers in the 5′ UTR, and transgene-derived transcripts were detected with primers spanning transgene-specific sequence immediately at the 5′ end of the Myc tag. Endpoint PCR products were resolved on 1% agarose gels and quantified by densitometry to calculate Ex/In ratios; a decrease indicates reduced exitron exclusion.

Functional nuclear splicing competence was assessed using established TDP-43-dependent sentinel events, namely *STMN2* cryptic-exon inclusion and *POLDIP3* exon-3 skipping (Koide et al., 2026; Shiga et al., 2012). All qPCR reactions were performed in technical duplicates. Specificity was confirmed by melt-curve analysis, and amplification efficiency was verified for each primer pair. Primer sequences are provided in Supplementary Table 1.

### 2.8. Immunocytochemistry and imaging

For imaging of iPSC-derived neurons, cells were fixed in 4% paraformaldehyde, permeabilized with 0.1% Triton X-100, and blocked with 5% bovine serum albumin. Primary antibodies were anti-TDP-43 (Proteintech, 12892-1-AP), anti-Myc (9B11; Cell Signaling Technology, 2276S), and anti-TUJ1 (Abcam, ab7751). Images were acquired on a Keyence BZ-X810. Exposure settings were held constant across each experiment and verified to remain within the linear dynamic range.

Nuclei were segmented based on Hoechst staining. Cytoplasmic mean intensities were extracted from somatic regions of interest (ROIs) defined within the TUJ1-positive cell body and outside the nuclear mask. The nuclear-to-cytoplasmic ratio (N/C) was calculated from these ROIs and log_2_-transformed for analysis. Cytoplasmic TDP-43 foci were defined as discrete, non-nuclear TDP-43-immunoreactive structures with signal intensity clearly above the surrounding diffuse cytoplasmic signal under identical acquisition settings.

### 2.9. Structural prediction

To assess whether the engineered modifications altered the N-terminal dimerization interface, full-length TDP-43 sequences (residues 1–414), with or without the N-terminal NES insertion or the ΔNLS mutations, were submitted to AlphaFold 3 (Abramson et al., 2024). Predicted dimeric assemblies (WT homodimer, WT/+NES heterodimer, and WT/ΔNLS heterodimer) were inspected comparatively and visualized in UCSF ChimeraX (v1.10.1) (Pettersen et al., 2021).

### 2.10. Statistics

Statistical analyses were performed in R (v4.4.1). Two-group comparisons used Student’s *t*-test, Welch’s *t*-test, or paired *t*-test, as appropriate. For comparisons of multiple treatment groups against a control, one-way ANOVA followed by Dunnett’s post hoc test was used. Correlations were assessed using Pearson’s correlation coefficient or Spearman’s rank correlation. For per-cell imaging data, linear mixed-effects models (LMMs) were fitted using the lmerTest package, with experimental group as a fixed effect and image field as a random intercept to account for non-independence of cells acquired within the same field. Unless otherwise specified, *n* denotes the number of biological replicates. Data are presented as mean ± SD unless otherwise indicated. All tests were two-sided, and *P* < 0.05 was considered statistically significant.

## 3. Results

### 3.1. NES–TDP-43 preserves N-terminal domain (NTD)-mediated dimerization and redistributes co-expressed WT TDP-43 toward the cytoplasm

To determine whether appending an N-terminal NES perturbs the NTD-mediated dimerization interface of TDP-43 (Afroz et al., 2017; Shiina et al., 2010; Zhang et al., 2013), we quantified hetero-oligomerization between EGFP–TDP-43 (bait) and Myc-tagged variants (Fig. 1A). In cytoplasmic GFP-Trap co-immunoprecipitation assays, +NES co-recovered with EGFP–TDP-43 at WT-comparable levels (*P =* 0.89), whereas ΔNLS association was reduced by 0.77 log_2_ units relative to WT (*P =* 0.014; Fig. 1B, C). RNase A and RNase T1 treatment did not materially diminish Co-IP recovery, indicating that the interaction is predominantly protein-mediated rather than RNA-bridged (Supplementary Fig. S1). These biochemical findings were consistent with AlphaFold 3 predictions: WT/WT and WT/+NES dimers were compatible with the canonical NTD–NTD interface, while the WT/ΔNLS model showed interface distortion that paralleled the reduced Co-IP recovery (Supplementary Fig. S2).

We then asked whether co-recruitment by export-biased TDP-43 redistributes the nuclear pool. In fixed-cell imaging, Myc-NES–TDP-43 was enriched in the cytoplasm and was accompanied by cytoplasmic redistribution of EGFP–TDP-43 (Δlog_2_(N/C) = −0.80; *P =* 0.005), whereas ΔNLS produced no significant shift (Δlog_2_(N/C) = +0.21; *P =* 0.63) (Fig. 1D, E). Within the +NES group, EGFP–TDP-43 log_2_(N/C) decreased as the Myc-NES/EGFP ratio rose (slope β = −0.36 ± 0.06 log_2_(N/C) per unit log_2_(Myc/EGFP); *P =* 3.97 × 10^−7^; Fig. 1F), pointing to dose-dependent co-recruitment of nuclear TDP-43 by the export-biased construct. WT showed no such relationship (β = 0.08 ± 0.17; *P =* 0.65). Together, these findings indicate that Myc-NES–TDP-43 preserves NTD-mediated dimerization capacity and redistributes co-expressed WT TDP-43 toward the cytoplasm.

### 3.2. Low-load NES–TDP-43 partitions disproportionately into the cytoplasm and detergent-insoluble fraction

To track how a modest imposed export bias propagates through TDP-43 homeostasis, we generated a single-clone Tet-On HEK293T line expressing *Myc-NES–TDP-43–3′ UTR* (Fig. 2A). Doxycycline titration produced a dose-dependent rise in whole-cell RIPA-soluble Myc-NES–TDP-43 (Pearson *r* = 0.79, *n* = 10, *P =* 0.006; Fig. 2B). All subsequent experiments used 1.0 µg mL^−1^ doxycycline, which held whole-cell RIPA-soluble Myc-NES well below endogenous TDP-43 (Myc-NES/endogenous ≈ 0.08; Fig. 2B), inside a strict low-load window.

**Figure 2.**
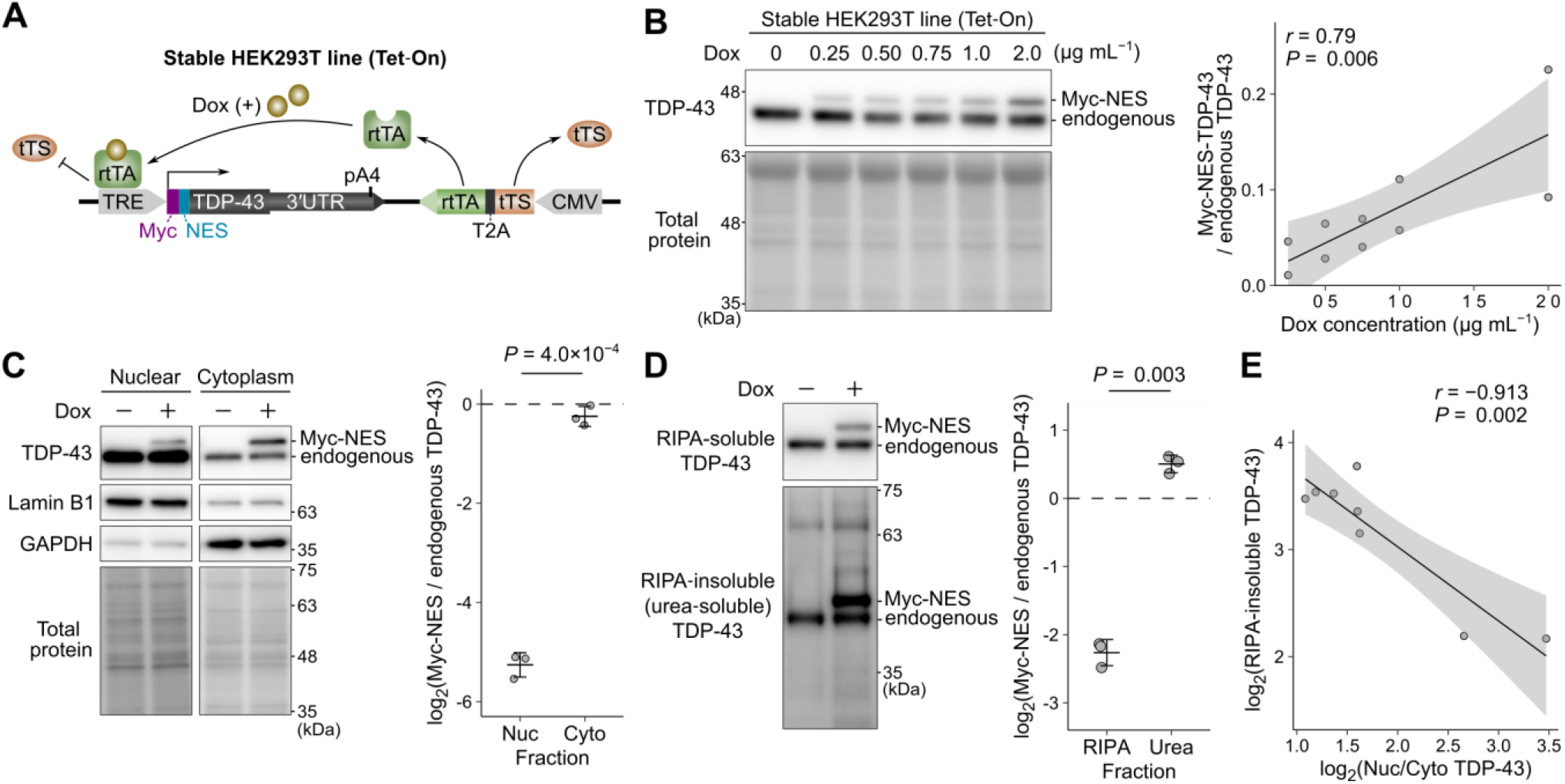
Low-load inducible NES–TDP-43 shows disproportionate cytoplasmic and detergent-insoluble partitioning. (A) Schematic of the Tet-On HEK293T line expressing *Myc-NES–TDP-43–3′ UTR*. (B) Doxycycline (Dox) titration (whole-cell RIPA-soluble fraction). Left: immunoblot of endogenous and Myc-NES bands (total-protein loading control). Right: dose-response of Myc-NES/endogenous TDP-43 ratios (linear fit ± 95% CI; *n* = 10). (C) Nucleocytoplasmic fractionation (1.0 µg mL^−1^ Dox). Left: immunoblots confirming fractionation purity (Lamin B1, nuclear; GAPDH, cytoplasmic). Right: relative abundance of Myc-NES in cytoplasmic versus nuclear fractions (paired *t*-test on log_2_ ratios; *n* = 3). (D) Solubility profiling. Immunoblots of paired RIPA-soluble and RIPA-insoluble (urea-soluble) fractions. Right: preferential partitioning of Myc-NES into the RIPA-insoluble fraction (paired *t*-test on log_2_ ratios; *n* = 3). (E) Relationship between localization and insolubility. Total TDP-43 N/C ratio [log_2_] plotted against total RIPA-insoluble burden [log_2_] (Pearson correlation with linear fit ± 95% CI; *n* = 8). Data represent mean ± SD.

Despite this low whole-cell soluble abundance, the protein was distributed asymmetrically between compartments. Relative to endogenous TDP-43 in matched fractions of uninduced controls, Myc-NES–TDP-43 reached 2.6 ± 0.4% of endogenous in the nuclear fraction but 84.8 ± 12.1% in the cytoplasmic fraction (Fig. 2C)—about a 32.6-fold cytoplasmic-over-nuclear enrichment (paired comparison of fractional representation, *P =* 4.0 × 10^−4^). Because the baseline cytoplasmic pool is small, even modest amounts of export-biased protein produce a disproportionate cytoplasmic increase.

Solubility was skewed in the same direction. Under the same induction, the Myc-NES/endogenous TDP-43 ratio was 0.210 ± 0.027 in the RIPA-soluble fraction but 1.42 ± 0.12 in the RIPA-insoluble (urea-soluble) fraction—a 6.8-fold higher insoluble-to-soluble ratio than for endogenous TDP-43 (*P =* 0.003; Fig. 2D). Across conditions, the total TDP-43 N/C ratio (endogenous + exogenous, including fragments) was inversely correlated with total RIPA-insoluble burden (Pearson *r* = −0.913, *n* = 8, *P =* 0.002; Fig. 2E, Supplementary Fig. S3).

### 3.3. Endogenous *TARDBP* knockdown relieves 3′ UTR-coupled autorepression under fixed export bias but does not restore the soluble TDP-43 pool

How does *TARDBP* autoregulatory feedback respond when endogenous *TARDBP* is knocked down while export bias is held constant? Using an inducible HEK293T line expressing *Myc-NES–TDP-43–3′ UTR* from a construct bearing the *TARDBP* 3′ UTR, we kept doxycycline induction constant and selectively depleted endogenous *TARDBP* with a 5′ UTR-targeting siRNA (si*TARDBP*) (Fig. 3A). Because the transgene lacks the native 5′ UTR, si*TARDBP* preferentially reduces the endogenous *TARDBP* while sparing the export-biased transgene.

**Figure 3.**
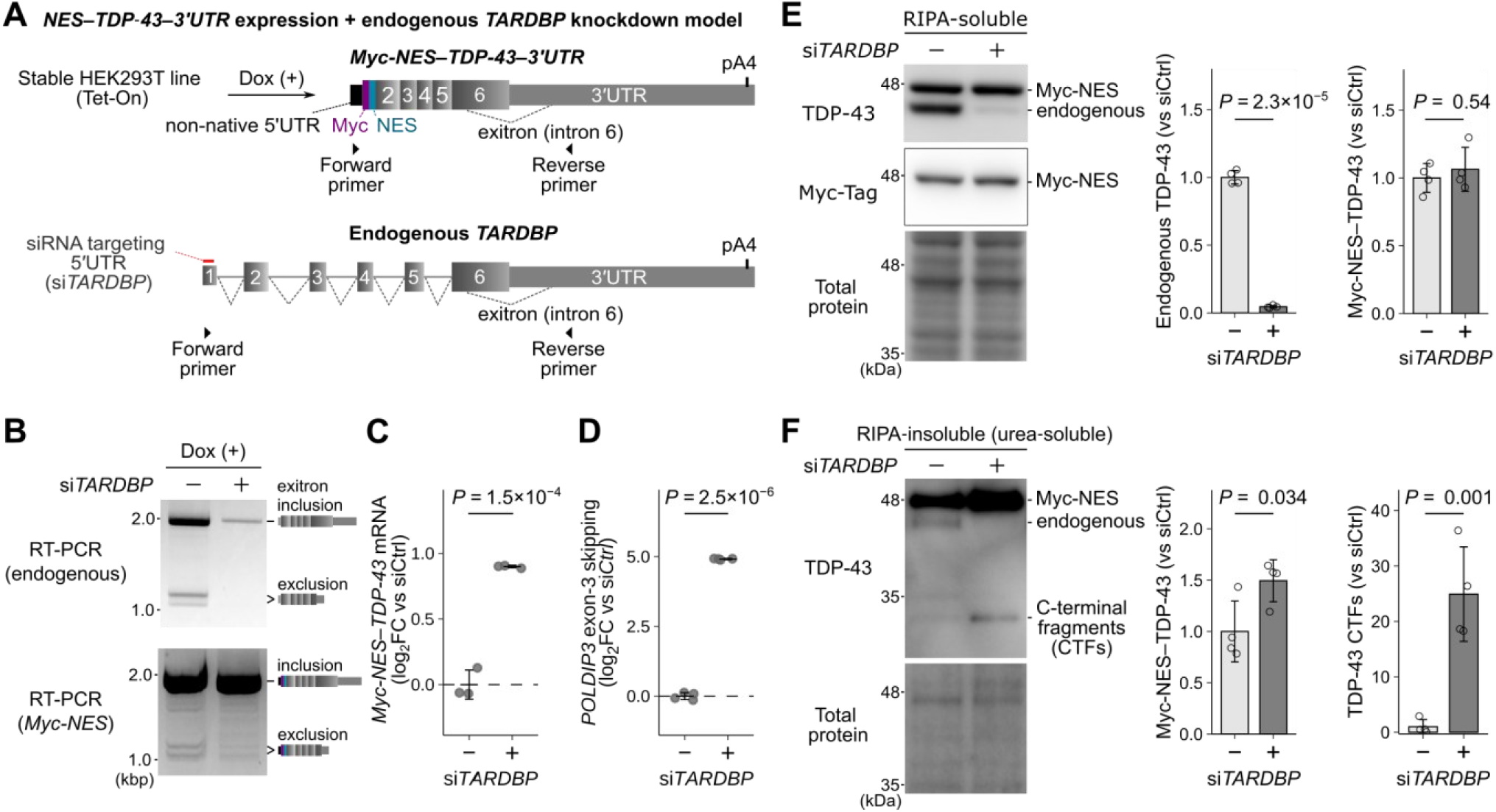
Endogenous *TARDBP* knockdown relieves 3′ UTR-coupled autorepression under fixed export bias but does not restore the soluble TDP-43 pool. (A) Schematic. Endogenous *TARDBP* was depleted using a 5′ UTR-targeting siRNA under fixed Myc-NES induction; primer sites for source-specific exitron analysis are indicated. (B) Source-specific endpoint RT-PCR showing reduced exitron exclusion in *Myc-NES– TDP-43–3′ UTR* transcripts following endogenous depletion. (C) *Myc-NES–TDP-43* transcript abundance normalized to *RPLP1*. RT-qPCR quantification showing upregulation upon weakened autorepression (log_2_ fold-change vs. siCtrl). (D) Nuclear function readout. *POLDIP3* exon-3 skipping indicates persistent nuclear loss of function. (E) RIPA-soluble protein pools. Immunoblots and quantification of endogenous and Myc-NES bands (total-protein normalization). Note stable soluble Myc-NES levels despite increased transcript abundance. (F) RIPA-insoluble (urea-soluble) burden. Immunoblots of full-length and C-terminal fragments (CTFs) with corresponding quantification. Mean ± SD; *n* = 3–4; two-sided *t*-tests.

Source-specific endpoint RT-PCR showed a marked reduction in the exitron-excluded Myc-NES band after si*TARDBP* (Fig. 3B), consistent with reduced exitron exclusion and relief of 3′ UTR-coupled autorepression. RT-qPCR detected a concomitant increase in total transgene-derived *TARDBP* mRNA (1.87 ± 0.01-fold vs. siCtrl; *P =* 1.5 × 10^−4^; Fig. 3C), consistent with preferential accumulation of the more stable exitron-included transcript species when autorepression is relieved.

The transcript increase did not translate into expansion of the soluble TDP-43 pool or recovery of the soluble TDP-43-dependent splicing. *POLDIP3* exon-3 skipping increased (*P* = 2.5 × 10^−6^; Fig. 3D), indicating persistent nuclear loss of function. In the RIPA-soluble fraction, Myc-NES–TDP-43 protein was unchanged (*P* = 0.54), whereas endogenous TDP-43 fell as expected after si*TARDBP* (*P* = 2.3 × 10^−5^; Fig. 3E). Solubility profiling resolved the disconnect: in the RIPA-insoluble (urea-soluble) fraction, full-length Myc-NES–TDP-43 increased (*P =* 0.034), and C-terminal fragments more so (*P =* 0.001; Fig. 3F). Thus, with export bias held constant, relief of *TARDBP* mRNA autorepression increases transcript supply, but the additional protein partitions into insoluble and fragmented species rather than replenishing the soluble, splice-competent TDP-43 pool.

### 3.4. Increasing NES–TDP-43 load drives progressive nuclear dysfunction in neuron-like cells despite soluble TDP-43 restoration

The next question was whether persistent export bias in a neuronal context produces graded nuclear dysfunction within the low-load range, and whether restoring soluble abundance can reverse it. Differentiated SH-SY5Y cells were transduced with lentiviral *Myc-NES–TDP-43–3′ UTR* and analyzed 7 days later (Fig. 4A). Varying viral input gave a graded increase in whole-cell RIPA-soluble Myc-NES–TDP-43, reaching about 30% of endogenous TDP-43 at the upper bound of the low-load range (Fig. 4B).

**Figure 4.**
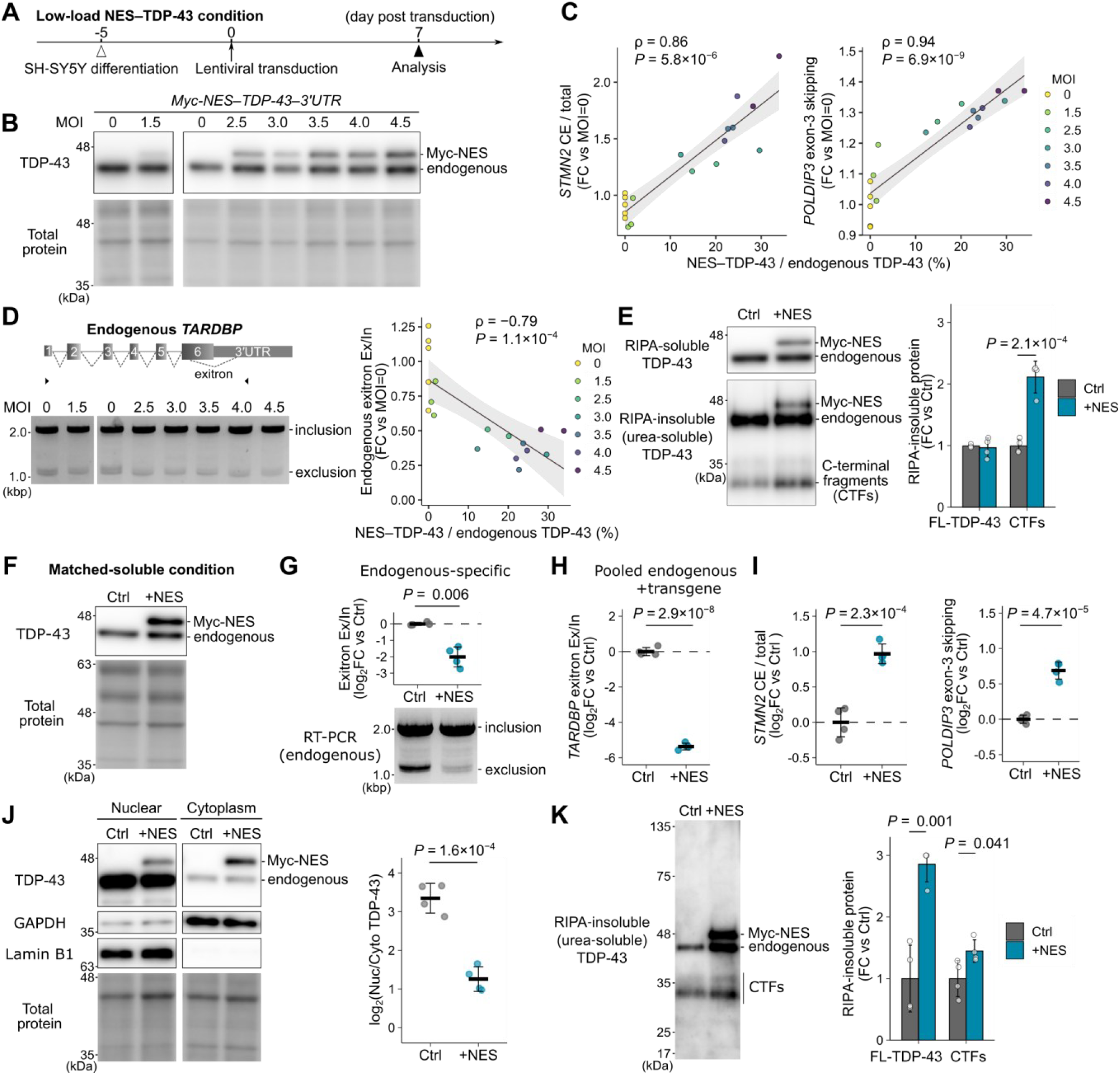
Increasing NES–TDP-43 load drives progressive nuclear dysfunction in neuron-like cells despite soluble TDP-43 restoration. (A–E) Low-load multiplicity of infection (MOI) series analyzed 7 days after lentiviral transduction of differentiated SH-SY5Y cells with *Myc-NES–TDP-43–3′ UTR*. (A) Experimental timeline. (B) Immunoblots showing graded RIPA-soluble Myc-NES–TDP-43 expression across MOIs. The immunoblots for MOI 0, 1.5 and MOI 0/2.5–4.5 were performed on different membranes. (C) *STMN2* cryptic-exon (CE) inclusion (left) and *POLDIP3* exon-3 skipping (right) plotted against soluble NES– TDP-43/endogenous TDP-43 ratios. Correlations were assessed by Spearman’s rank correlation. (D) RT-PCR products for MOI 0, 1.5 and MOI 0/2.5–4.5 were resolved on separate gels. Endogenous *TARDBP* exitron exclusion-to-inclusion ratio (Ex/In) plotted against soluble NES–TDP-43/endogenous TDP-43 ratios. (E) Paired RIPA-soluble and RIPA-insoluble (urea-soluble) fractions at MOI 4.0 showing insoluble accumulation of C-terminal fragments (CTFs). (F–K) Soluble-matched condition analyzed at day 7. (F) Immunoblot confirming soluble Myc-NES–TDP-43 abundance comparable to or exceeding endogenous levels. (G, H) Persistence of exitron-processing defects measured by endogenous-specific RT-PCR (G) and pooled RT-qPCR (H). (I) *STMN2* cryptic-exon inclusion and *POLDIP3* exon-3 skipping persist despite restoration of soluble abundance. (J) Nucleocytoplasmic fractionation showing reduced total TDP-43 N/C ratio. (K) Solubility profiling showing increased RIPA-insoluble (urea-soluble) burden. Data represent mean ± SD.

Nuclear TDP-43 readouts deteriorated progressively across this range. *STMN2* cryptic-exon inclusion increased steeply with the soluble NES–TDP-43/endogenous TDP-43 ratio (Spearman ρ = 0.86, *P* = 5.8 × 10^−6^; Fig. 4C, left). *POLDIP3* exon-3 skipping also tracked NES–TDP-43 load: the correlation was tighter (ρ = 0.94, *P* = 6.9 × 10^−9^) but the absolute fold-changes were smaller than for *STMN2* (Fig. 4C, right). Endogenous *TARDBP* exitron processing shifted toward inclusion, reflected as a decrease in Ex/In with rising soluble NES–TDP-43 (ρ = −0.79, *P* = 1.1 × 10^−4^; Fig. 4D). Even when soluble exogenous TDP-43 stayed well below endogenous levels, increasing export-biased load was therefore associated with progressively abnormal nuclear splicing readouts.

To determine whether biochemical conversion had already begun within this regime, we examined paired RIPA-soluble and RIPA-insoluble (urea-soluble) fractions at MOI 4.0, near the upper bound of the low-load window. Myc-NES–TDP-43 was readily detected in the insoluble fraction together with increased fragments (*P =* 2.1 × 10^−4^; Fig. 4E); insoluble accumulation therefore co-occurred with the exitron and splicing abnormalities seen across this range.

We then asked whether expanding the soluble pool could normalize these readouts while cytoplasmic bias remained. Higher viral input produced a soluble-matched condition in which RIPA-soluble Myc-NES–TDP-43 reached or exceeded endogenous abundance (Fig. 4F); nuclear readouts nevertheless remained abnormal relative to control. Endogenous-specific RT-PCR again showed marked loss of the exitron-excluded band in +NES cultures (Fig. 4G), and pooled *TARDBP* RT-qPCR (endogenous + exogenous) showed a decrease in log_2_(Ex/In) (*P =* 2.9 × 10^−8^; Fig. 4H). Isoform-level deconvolution attributed the pooled shift to reduced abundance of the major exitron-excluded isoforms (Supplementary Fig. S4). Because the splice-competent transgene also contributes to those isoforms, the net decrease most likely reflects fewer correctly processed endogenous *TARDBP* transcripts. Functionally, *STMN2* cryptic-exon inclusion and *POLDIP3* exon-3 skipping persisted despite the restored soluble abundance (*P =* 2.3 × 10^−4^ and *P =* 4.7 × 10^−5^, respectively; Fig. 4I).

To understand why soluble-pool restoration failed to normalize function, we assessed compartmentalization and biochemical state. Nucleocytoplasmic fractionation revealed pronounced redistribution to the cytoplasm: total TDP-43 N/C fell to 23.3 ± 5.3% of control values, a 4.3-fold decrease (*P =* 1.6 × 10^−4^; Fig. 4J). Solubility profiling showed a parallel diversion into detergent-insoluble states, with full-length TDP-43 and fragments both increased (*P =* 0.001 and *P =* 0.041, respectively; Fig. 4K). Thus, in neuron-like cells, increasing NES– TDP-43 load drives progressive nuclear splicing dysfunction within the low-load range, and restoring soluble abundance alone is insufficient to rescue function while redistribution and insoluble partitioning persist.

### 3.5. iPSC-derived neurons recapitulate cytoplasmic redistribution and nuclear splicing defects under export bias

Finally, we tested whether the same imposed-cytoplasmic-bias / nuclear-dysfunction relationship is reproduced in human neurons. Human iPSC-derived neurons were transduced at DIV14 with Myc-NES–TDP-43–3′ UTR and analyzed at DIV21 (Fig. 5A). Under these conditions, RIPA-soluble exogenous Myc-NES was about 3.8% of endogenous TDP-43 (Fig. 5B), well within the low-load window defined in whole-cell soluble lysates.

**Figure 5.**
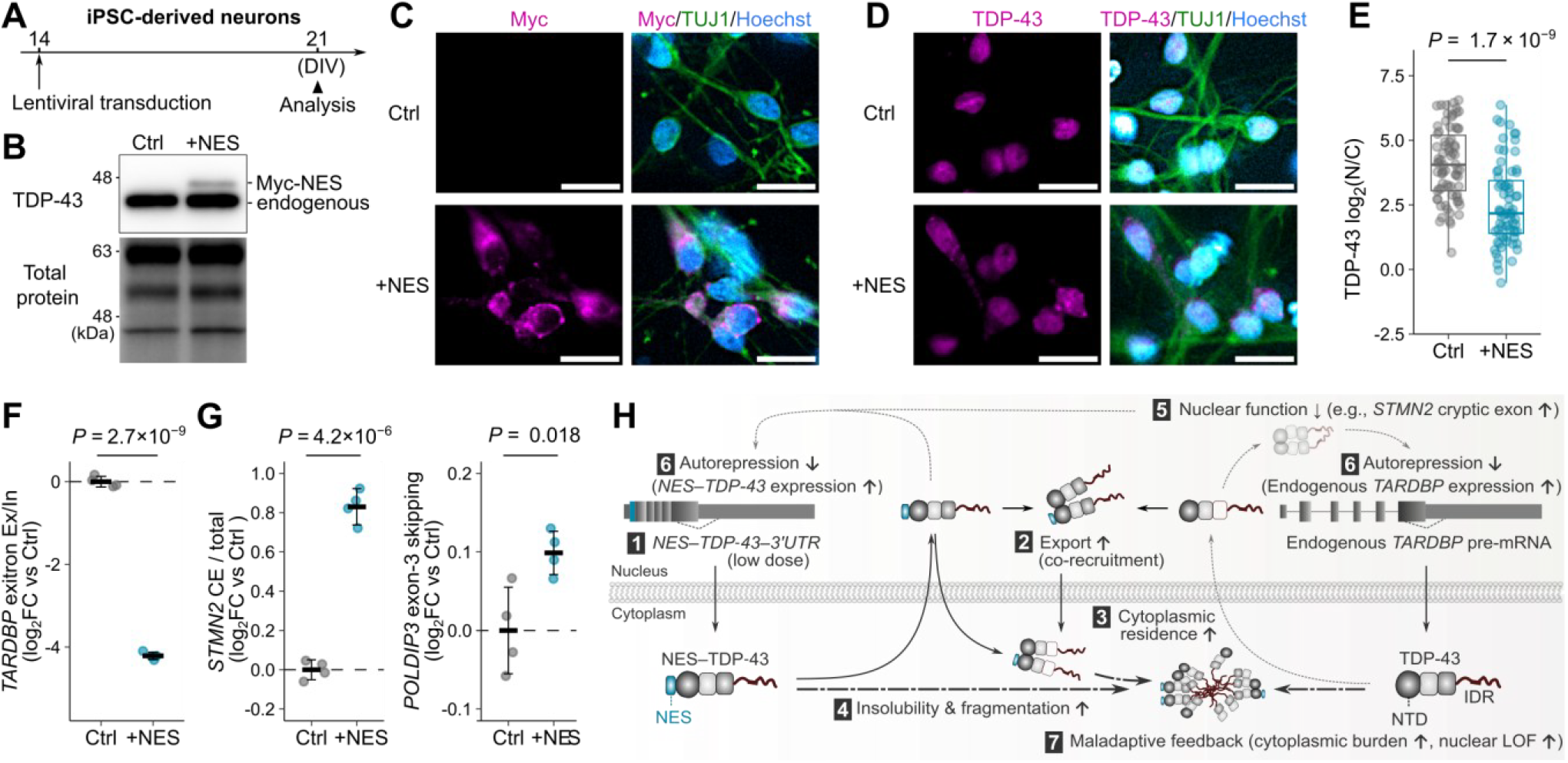
iPSC-derived neurons recapitulate cytoplasmic redistribution and nuclear splicing defects under export bias. (A) Timeline. Human iPSC-derived neurons were transduced at DIV14 and analyzed at DIV21. (B) Low-load verification. Immunoblot showing low Myc-NES abundance relative to endogenous TDP-43. (C–E) Representative immunocytochemistry showing Myc-tagged construct expression (C) and redistribution of total TDP-43 toward the cytoplasm (D). Scale bars, 20 µm. (E) Quantification showing a significant shift in TDP-43 log_2_(N/C) relative to control (LMM; image field as random intercept, *n* = 145 cells from 20 image fields). (F) Autoregulatory readout. Reduced pooled exitron exclusion-to-inclusion ratio [log_2_(Ex/In)] indicates altered autoregulatory processing consistent with reduced nuclear TDP-43 function. (G) Functional sentinels. Mis-splicing of *STMN2* and *POLDIP3* indicates nuclear loss-of-function signatures. Mean ± SD; *n* = 4; two-sided *t*-test (F, G). (H) Working model. (1–3) Export bias promotes NTD-mediated co-recruitment and cytoplasmic redistribution. (4) Cytoplasmic TDP-43 shows increased partitioning into detergent-insoluble and fragmented species. (5–6) Reduced nuclear TDP-43 is accompanied by splicing defects and weakening of 3′ UTR-coupled autorepression, increasing *TARDBP*/TDP-43 synthesis. (7) Under persistent cytoplasmic bias, this compensatory output may be diverted into insoluble or fragmented cytoplasmic species rather than restoring the nuclear functional pool. Thick dashed arrows denote reinforced flux; thin dotted arrows denote attenuated processes.

Even so, total TDP-43 redistributed toward the cytoplasm. Quantification of TDP-43 immunofluorescence showed a significant decrease in log_2_(N/C) relative to control (*P =* 1.7 × 10^−9^), about a threefold reduction in the nuclear-to-cytoplasmic ratio (Fig. 5C–E). In keeping with the cytoplasmic excess, 42.1 ± 7.9% of Myc-NES–positive neurons carried discrete cytoplasmic TDP-43-positive foci.

Readouts of nuclear TDP-43 function tracked the redistribution. RT-qPCR profiling of *TARDBP* exitron in pooled endogenous plus transgene-derived transcripts showed a significant reduction in log_2_(Ex/In) (*P =* 2.7 × 10^−9^; Fig. 5F), and isoform-level deconvolution revealed both increased exitron inclusion and reduced abundance of the major exitron-excluded isoforms (Supplementary Fig. S5). *STMN2* cryptic-exon inclusion and *POLDIP3* exon-3 skipping were both increased (*P =* 4.2 × 10^−6^ and *P =* 0.018, respectively; Fig. 5G). Modest export bias was therefore enough to redistribute TDP-43 toward the cytoplasm and to produce autoregulatory and splicing signatures of nuclear dysfunction in human iPSC-derived neurons (Fig. 5H).

## 4. Discussion

This study asked whether native 3′ UTR-coupled autoregulation remains protective when TDP-43 nucleocytoplasmic balance is persistently shifted toward the cytoplasm. Our data indicate that it does not. In differentiated SH-SY5Y cells within the low-load range, export-biased TDP-43 produced progressive abnormalities in exitron processing and TDP-43-dependent splicing, together with insoluble accumulation, and these abnormalities persisted even when soluble NES–TDP-43 was raised to endogenous-equivalent levels. Human iPSC-derived neurons likewise showed cytoplasmic redistribution, discrete TDP-43-positive foci, and splicing changes indicative of reduced nuclear TDP-43 function. When autorepression was weakened by selective depletion of endogenous TDP-43, the resulting rise in transgene-derived *TARDBP* transcripts did not enlarge the soluble functional pool; the extra protein appeared mainly in insoluble and fragmented fractions. We therefore propose that persistent cytoplasmic bias uncouples compensatory *TARDBP* upregulation from restoration of the functional nuclear pool.

Several limitations of this experimental framework should be considered when interpreting these findings. First, the PKIα NES was used as an engineered perturbation to impose persistent cytoplasmic bias; it is not intended to represent the native export route of TDP-43 or to model any single upstream lesion in ALS/FTLD. In disease, altered TDP-43 partitioning may arise through impaired nuclear import, nuclear pore dysfunction, or stress-associated cytoplasmic retention (Chou et al., 2018; Combe et al., 2026; Gasset-Rosa et al., 2019; Prpar Mihevc et al., 2017; Woerner et al., 2016; Zhang et al., 2018). Second, the low-load definition used here is based on whole-cell RIPA-soluble exogenous TDP-43 and does not imply compartment-specific or local cytoplasmic concentrations. Third, detergent insolubility, fragmentation, and discrete cytoplasmic foci are operational signatures of biochemical conversion, but they do not, on their own, establish the ultrastructure, phospho-epitope composition, or maturation state of disease inclusions (Arseni et al., 2022; Hasegawa et al., 2008; Mori et al., 2008). The agreement across HEK293T cells, differentiated SH-SY5Y cells, and iPSC-derived neurons supports the robustness of the observed coupling, but none of these systems reproduces the multicellular and temporal context of ALS/FTLD. We also do not directly resolve the kinetics by which newly synthesized TDP-43 is partitioned between productive nuclear and non-productive cytoplasmic states; that question requires live, isoform-resolved tracking that is beyond the reach of the present design.

These results identify a specific failure mode for TDP-43 homeostasis under native 3′ UTR-coupled control: when the cytoplasmic shift does not resolve, the autoregulatory circuit can keep producing transcript without replenishing the functional nuclear pool, and the surplus protein is exposed to the same cytoplasmic environment that drives insolubility and fragmentation. A protective buffering circuit, in other words, becomes maladaptive precisely because its substrate is mislocalized at the time of compensation. The export-biased system described here provides a tractable model for studying that early homeostatic failure under native autoregulatory control.

## Declarations

### Availability of data and material

The datasets generated and/or analyzed during the current study are available in the Zenodo repository (DOI: https://doi.org/10.5281/zenodo.19968143). Reviewer access links: https://zenodo.org/records/19968143?preview=1&token=eyJhbGciOiJIUzUxMiJ9.eyJpZCI6IjcyNmViZDY3LTRmMzItNDg2Mi04N2FhLTE0YzVlNGIwZGFkOCIsImRhdGEiOnt9LCJyYW5kb20iOiIyYmZkODJkNmNlYTU2Y2UxMTZhZTRkYWQ3ODUzZmUyMyJ9.o_VI6mdbOyvD2aiVBJryXouQfVeKfuzk6czR3iJWspFTPA2TEW-KbYKFn8-S-hSLxkoN2XM2Cg47CiTfxA4Y_w

### Competing interests

A.S. and O.O. are inventors on WO2022113799A1, which covers antisense oligonucleotides targeting the *TARDBP* exitron; this patent has been granted in Japan and the United States, with a European application currently under examination. A.S., T.Y. and O.O. are inventors on WO2023204313A1, which covers the aggregation-suppressive function of IDRsTDP. A.S., S.K. and O.O. are inventors on WO2025234407A1 and WO2026009728A1, which cover additional antisense oligonucleotide designs targeting the *TARDBP* exitron; these applications are pending. All patent applications are assigned to Niigata University. The remaining authors declare no competing interests.

### CRediT authorship contribution statement

**Genri Toyama:** Conceptualization, Investigation, Formal analysis, Writing. **Shingo Koide:** Conceptualization, Investigation, Formal analysis. **Takuma Yamagishi:** Conceptualization, Formal analysis. **Aya Washida:** Investigation. **Ryutaro Hanyu:** Investigation. **Osamu Onodera:** Writing, Supervision. **Akihiro Sugai:** Conceptualization, Investigation, Formal analysis, Writing, Funding acquisition.

## Acknowledgements

This work was supported by the Japan Agency for Medical Research and Development (AMED) (JP22ek0109603; AS), Japan Society for the Promotion of Science (JSPS) KAKENHI (JP24K22094 and JP25K02462; AS), and New Sustainable Growth (NSG) group, which supports the Advanced Treatment of Neurological Diseases Branch, Brain Research Institute, Niigata University (AS).

## Declaration of generative AI and AI-assisted technologies in the manuscript preparation process

During the preparation of this work the authors used ChatGPT (OpenAI) in order to rephrase English sentences and verify English grammar. After using this tool/service, the authors reviewed and edited the content as needed and take full responsibility for the content of the published article.

---

**Supplementary Table 1.**
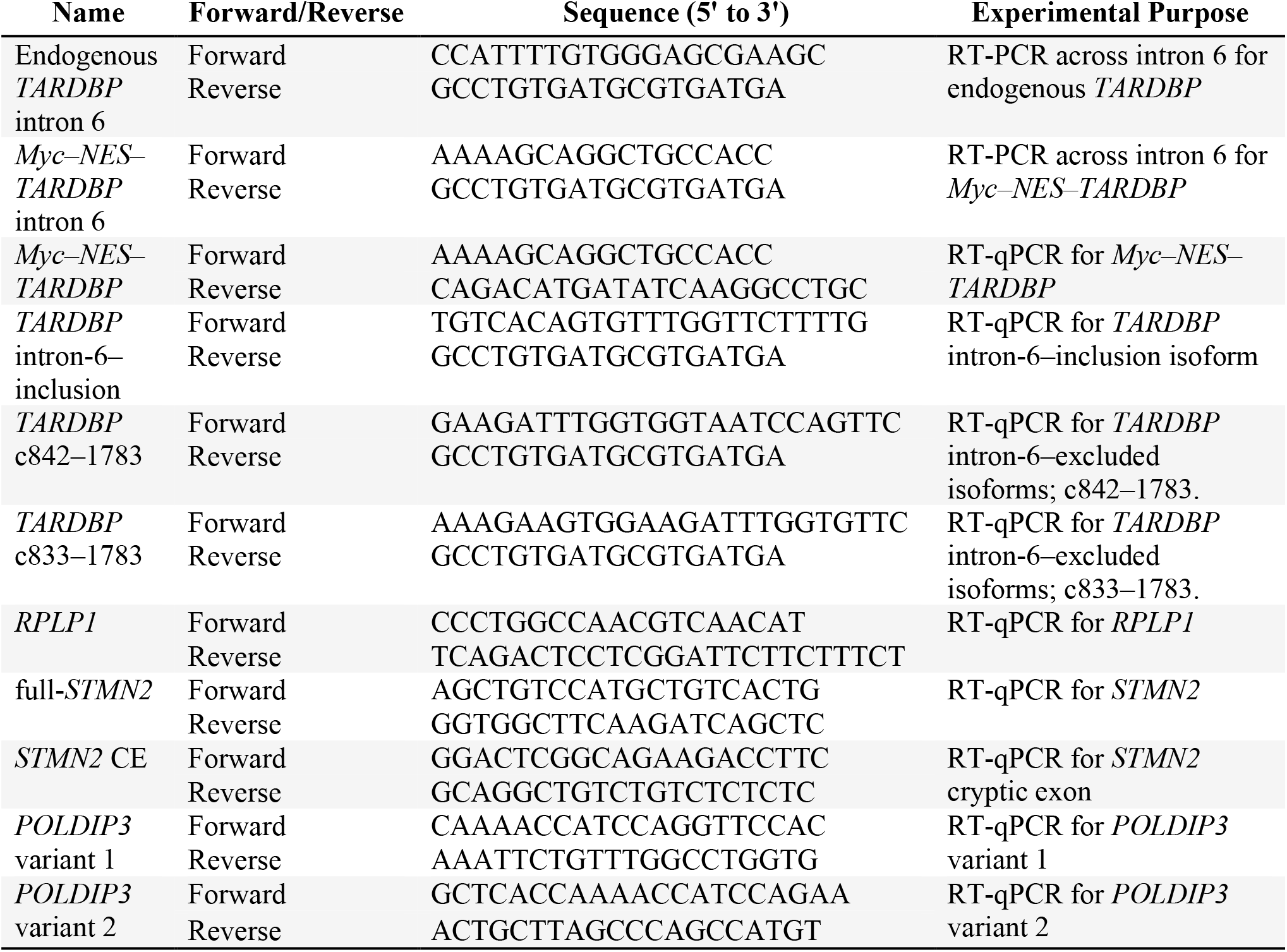
Primer sequences.

## Notes

### Summary of Updates

The manuscript has been thoroughly revised throughout, with new data added to Figure 4.

