## Supplemental Figure for "Export‑biased, 3′UTR‑preserving TDP‑43 model links nuclear loss to cytoplasmic aggregation in ALS/FTLD"

### Figure S1

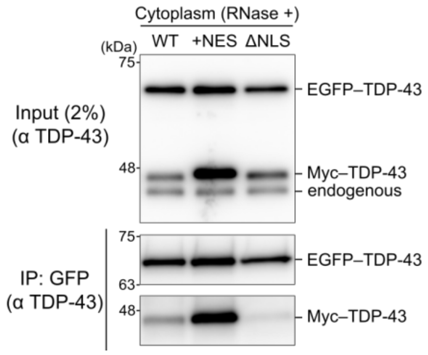


**Supplementary Figure S1. Cytoplasmic association of +NES–TDP-43 is largely RNA-independent.** Cytoplasmic GFP-Trap Co-IPs performed after RNase A/T1 digestion. Immunoblots detect EGFP–TDP-43 (bait), Myc–TDP-43 (prey), and endogenous TDP-43. Co-recovery of WT and +NES persists after RNase treatment, whereas ΔNLS association remains reduced, supporting a predominantly protein-mediated rather than RNA-bridged interaction.

### Figure S2

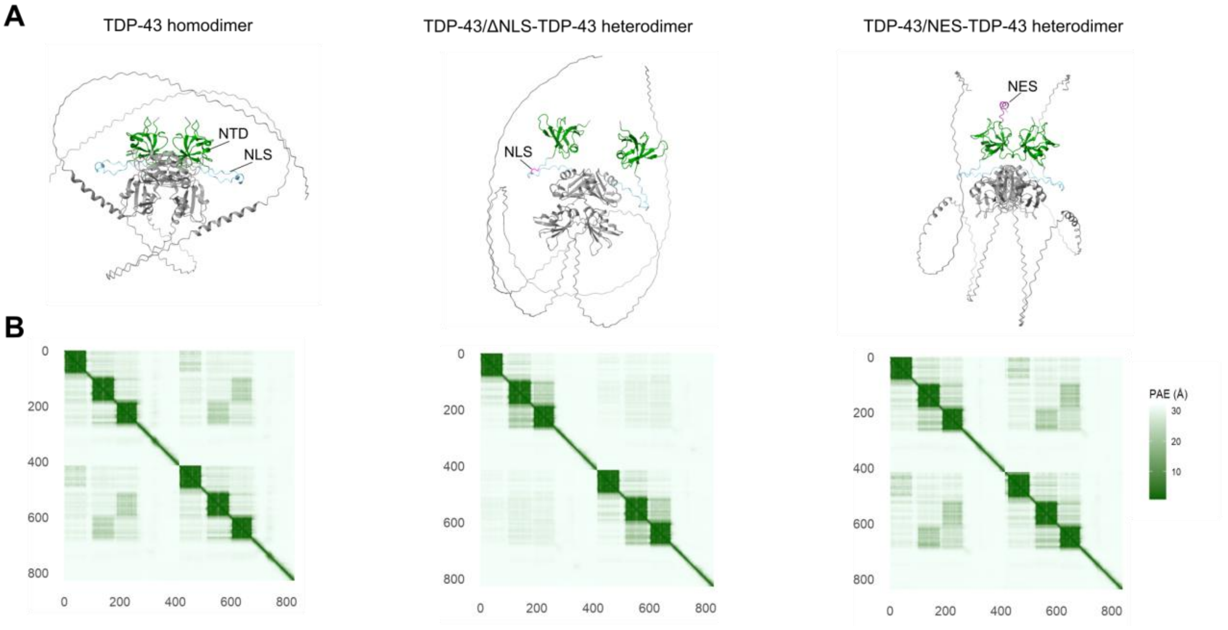


**Supplementary Figure S2. AlphaFold 3 predictions are compatible with preserved N-terminal dimerization of +NES–TDP-43.** (A) Structural predictions of N-terminal dimers: WT/WT homodimer (left), WT/ΔNLS heterodimer (middle), WT/+NES heterodimer (right). The canonical NTD–NTD head-to-head interface is preserved in WT and +NES models, suggesting that the flexible N-terminal NES extension does not sterically hinder assembly. In contrast, the WT/ΔNLS heterodimer shows a distorted, asymmetric interface geometry. (B) Predicted alignment error (PAE) maps. Low inter-chain PAE indicates a more confident structural interface for WT and +NES, whereas WT/ΔNLS shows elevated inter-chain error, consistent with a less stable interaction model.

### Figure S3

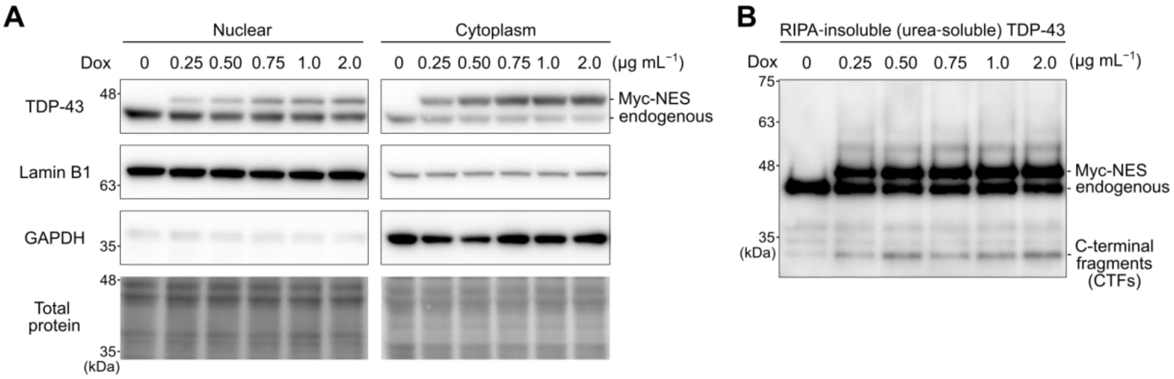


**Supplementary Figure S3. Increasing NES induction is associated with greater detergent-insoluble TDP-43 accumulation.** Source data for Fig. 2E. (A) Nucleocytoplasmic fractionation across the doxycycline range (0–2.0 µg $mL^{-1}$). (B) Matched RIPA-insoluble (urea-soluble) fractions showing dose-dependent accumulation of full-length Myc-NES, C-terminal fragments, and high-molecular-weight smears.

### Figure S4

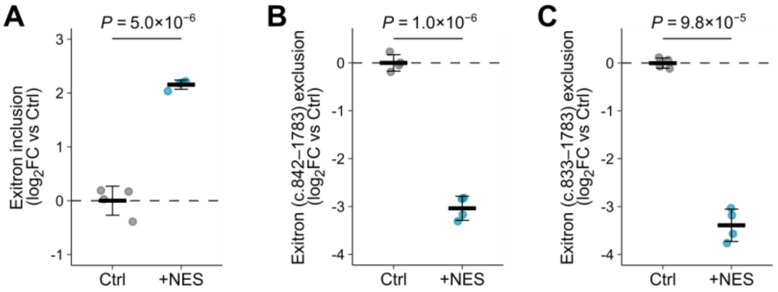


**Supplementary Figure S4. Deconvolution of *TARDBP* processing in SH-SY5Y cells supports a shift away from excluded isoforms despite expression of a splice-competent transgene.** Source data for Fig. 4H (SH-SY5Y, day 7, soluble-matched condition). RT-qPCR of pooled (endogenous + exogenous) transcripts normalized to *RPLP1*. (A) Increased exitron inclusion, consistent with weakening of 3′ UTR-coupled autorepression. (B, C) Decreased major exitron-excluded isoforms (c.842–1783 and c.833–1783). Because the transgene is splice-competent and contributes to the excluded pools, the net decrease supports a shift away from correctly excluded isoforms in the endogenous transcript population. Mean ± SD; $n = 4$; two-sided *t*-tests.

### Figure S5

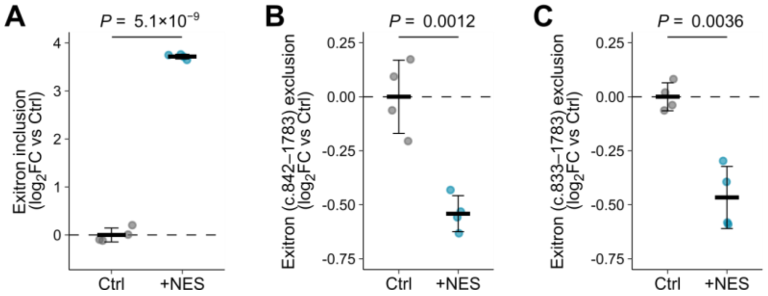


**Supplementary Figure S5. Deconvolution of *TARDBP* processing in human iPSC-derived neurons supports altered exitron processing despite expression of a splice-competent transgene.** Source data for Fig. 5F (iPSC-derived neurons, DIV21). RT-qPCR of pooled (endogenous + exogenous) transcripts normalized to *RPLP1*. (A) Increased exitron inclusion, consistent with weakening of 3′ UTR-coupled autorepression. (B, C) Decreased major exitron-excluded isoforms (c.842–1783 and c.833–1783). As in SH-SY5Y cells, the net decrease despite expression of splice-competent Myc-NES transcripts supports altered endogenous *TARDBP* processing that contributes to the reduced Ex/In ratio in Fig. 5F. Mean ± SD; $n = 4$; two-sided $t$-tests.
